## Supplementary Materials and Methods for "Reference-free cell-type deconvolution of multi-cellular pixel-resolution spatially resolved transcriptomics data"

**Table of Contents**

A. Supplementary Notes
   1. Suitability of ST data for LDA
   2. Selection of an appropriate *K*
   3. Bias gene selection in MERFISH benchmark

B. Supplementary Methods
   - Deconvolution of simulated MERFISH ST data with STdeconvolve
   - Deconvolution of simulated MERFISH ST data with supervised and semi-supervised reference-based deconvolution approaches
      - Deconvolution of simulated MERFISH ST data by reference-based deconvolution approaches using a reference with missing cell-types
      - Deconvolution of simulated MERFISH ST data by reference-based deconvolution approaches using a mouse brain scRNA-seq reference
   - Simulating ST data using mixtures of cells from scRNA-seq
      - Simulating age perturbed ST data
      - Deconvolution compared to clustering analysis of simulated cell-type mixtures
      - Simulating and deconvolving immune infiltrated vs excluded ST data
      - Simulating and deconvolving ST data with uniform cell-type distributions
   - Deconvolution of simulated MERFISH mouse coronal ST data with BayesSpace and STdeconvolve
   - Clustering analysis of ST data of the mouse olfactory bulb (MOB)
   - Deconvolution of ST data of the MOB
      - Comparison of MOB biological replicates

    - Deconvolution of ST data of the MOB by reference-based deconvolution approaches using a reference with missing cell-types
    - Identification of MOB cell-type putative marker genes
    - Deconvolution of ST data of the MOB by reference-based deconvolution approaches using a mouse brain scRNA-seq reference
    - Comparing deconvolution of the MOB between STdeconvolve, SPOTlight, RCTD, and spatialDWLS
    - Identification of VLMC cell-type putative marker genes
- Deconvolution of DBiT-seq data with STdeconvolve
- Deconvolution of Slide-seq data
    - With STdeconvolve
    - With RCTD
    - Comparison of STdeconvolve and RCTD
- Deconvolution of 10X Visium data with STdeconvolve
- Clustering analysis of breast cancer sections
- Deconvolution of ST data of breast cancer sections with STdeconvolve
    - Gene set enrichment analysis of deconvolved breast cancer cell-types
    - Enrichment of deconvolved cell-types in breast cancer pathological labels
- Accuracy of STdeconvolve with respect to ST dataset size

C. Supplementary Figures

Figure S1. Simulated MERFISH MPOA ST dataset and STdeconvolve predictions for 9 cell-types across all 12 bregmas.

Figure S2. Comparison of deconvolved cell-types from STdeconvolve to ground truth cell-types of simulated MERFISH MPOA ST data.

Figure S3. Cell-type predictions of supervised and semi-supervised deconvolution approaches across simulated MERFISH MPOA ST dataset pixels with different single-cell transcriptomics references.

Figure S4. STdeconvolve deconvolves simulated ST data of aged and young tissues.

Figure S5. Single-cell transcriptional clustering of MERFISH data of a coronal section of the mouse brain.

Figure S6. Additional of the MOB.

Figure S7. Stability of deconvolved cell-types across MOB ST data of biological replicates.

Figure S8. Correlations between deconvolved cell-types of the MOB ST dataset by STdeconvolve and supervised deconvolution approaches.

Figure S9. Effect of missing OEC cell cluster in MOB reference on supervised deconvolution approaches.

Figure S10. Misassignment of VLMC cluster in MOB ST data by supervised deconvolution approaches when trained with cortex scRNA-seq reference.

Figure S11. Comparison of STdeconvolve cell-types to ground truth cell-types in high resolution pixel MERFISH MPOA ST datasets.

Figure S12. STdeconvolve pixel proportion of select deconvolved cell-types in 10X Visium data of the mouse brain.

Figure S13. STdeconvolve pixel proportions and transcriptional profiles of select deconvolved cell-types in DBiT-seq data of an E11 mouse embryo lower tail section.

Figure S14. STdeconvolve predicted cell-type proportions of Purkinje neurons and Bergmann glia in Slide-seq data of the mouse cerebellum compared to RCTD.

Figure S15. Runtime and memory usage by STdeconvolve.

Figure S16. Deconvolution of the breast cancer ST data by STdeconvolve.

Figure S17. STdeconvolve cell-types X3 and X13 of the breast cancer ST data.

Figure S18. STdeconvolve cell-types X15 of the breast cancer ST data.

Figure S19. Accuracy of deconvolution by STdeconvolve based on the number of pixels in the input dataset.

Figure S20. Deconvolution failures.

Figure S21. Comparison of STdeconvolve cell-types to ground truth neuronal subtypes of simulated MERFISH MPOA ST data.

D. Supplementary Tables

Table S1. Significant GO terms for STdeconvolve cell-type 15 derived from the breast cancer ST dataset.

E. Supplementary References

### A. Supplementary Notes

#### 1. <u>Suitability of ST data for LDA</u>

STdeconvolve is built around LDA, which seeks to represent latent "topics", or cell-types, as ideally non-overlapping groups of co-expressed, or frequently co-occurring, genes in different pixels. In this manner, successful application of LDA in deconvolving latent cell-types relies on several key assumptions that can be reasonably met in ST data.

First, LDA is a generative probabilistic model for collections of discrete data and as such assumes input data as discrete counts. Current data from ST technologies incorporate unique molecular identifiers to achieve gene-level counts across spatially resolved pixels and is therefore consistent with this assumption.

Next, as LDA represents ST pixels as sparse representations of latent cell-types[1], it assumes that a few cell-types are present in each pixel. Given the micron-resolution pixel size in ST data and the average size of cells, we can generally assume that a single cell-type or a mixture of a few cell-types are present in each ST pixel. Further, we can assume that the proportional distribution of cell-types across pixels is heterogeneous within tissues profiled by ST such that some pixels will have more of one cell-type and other pixels will have more of another cell-type.

Likewise, as LDA represents latent cell-types as groups of co-expressed or frequently co-occurring genes, it assumes that there will be multiple genes informative of each underlying cell-type. While we can generally assume this to be true to transcriptionally distinct cell-types, this does mean that LDA, and subsequently STdeconvolve, would not be suitable for resolving cell subtypes that are defined by very subtle combinations of minimally upregulated or downregulated genes with respect to other cell subtypes. Likewise, LDA, and subsequently STdeconvolve, would not be able to differentiate between cell-types that cannot be distinguished through distinct differences in proportional gene expression such as volume or morphology.

Next, LDA works best with a large number of documents. Consistent with this assumption, ST data measures the gene expression profiles of hundreds to thousands of spatially resolved pixels. Further, LDA works best when there are more documents than topics, else each document may be assigned to a different topic. For ST data, we can generally assume that there are more pixels than cell-types.

These unique features of ST data contribute to the utility of LDA in deconvolving the proportional representation and transcriptomic profiles of latent cell-types.

### 2. Selection of an appropriate *K*

STdeconvolve requires the number of cell-types to be deconvolved (i.e., *K*), to be determined *a priori*. This can be determine based on prior biological knowledge of the ST dataset, or alternatively by using a data-driven approach. STdeconvolve provides several metrics to help users make this choice of *K* (Methods, ‘*Selection of LDA model with optimal number of cell-types*’). For each fitted model, STdeconvolve reports the number of deconvolved cell-types with a mean pixel proportion < 5% (as default), i.e., “rare” cell-types. This default threshold of 5% was chosen because we found that the deconvolution accuracy of rare cell-types with mean pixel proportions below 5% was poor by STdeconvolve as well as reference-based deconvolution approaches when assessing the simulated MERFISH MPOA ST data. Additionally, when simulating 100 $\mu m^2$ resolution pixels using the MERFISH MPOA data, the mean number of total cells per pixel was approximately 18, which means a single cell contributes on average approximately 6% of the pixel cell-type proportion (Supplementary Figure S11E). Thus, cell-types that are assigned to be more than 5% of a pixel cell-type proportion are likely to be attributable to at least one cell in the pixel, which agrees with previously suggested thresholds by other reference-based deconvolution approaches[2]. We note that this default threshold of 5% can be adjusted by users depending on the resolution and tissue of the ST dataset.

#### 3. Bias gene selection in MERFISH benchmark

ST data provides full transcriptome but multi-cellular pixel resolution spatially resolved transcriptomic profiling. As such, not all genes available in ST data may be informative of underlying transcriptionally distinct cell-types. STdeconvolve thus begins with gene selection as described in Methods section `*Selection of genes for LDA Model*`.

To simulate ST data for benchmarking, we aggregated single-cell resolution MERFISH data to pixel resolution as described in Methods section `*Simulating ST data from single-cell resolution spatially resolved MERFISH data*`. It should be noted, however, that MERFISH is a targeted transcriptome profiling technique rather than full transcriptome. For this MERFISH dataset, a specific panel of 135 genes previously chosen to optimally distinguish primarily between neuronal subtypes. As such, only 33 remaining genes were available to distinguish between all major cell-types[3]. Therefore, these chosen genes represent a specific subset of overdispersed genes rather than all overdispersed genes that are skewed towards distinguishing between neurons, even though many more overdispersed genes may exist for distinguishing microglia and pericytes, for example. In this manner, neuronal subtypes may appear more transcriptionally distinct than certain cell-types for which few markers were included. Indeed, when we applied STdeconvolve to the simulated MERFISH MPOA ST data limited to only these

135 genes, deconvolved cell-types such as cell-types X2 and X8 both matched to excitatory neurons while cell-types X4 and X7 both matched to inhibitory neurons. Given that excitatory and inhibitory neurons could previously be further subdivided into finer neuronal sub-types, we sought to evaluate whether these deconvolved cell-types that matched to the excitatory and inhibitory neurons could represent these additional finer neuronal sub-types. To test this hypothesis, we further partitioned the ground-truth excitatory and inhibitory cell-types into additional sub-types based on previous annotations, resulting in 76 total non-neuronal cell-types and neuronal subtypes[3]. Comparing the deconvolved transcriptional profiles of X2, X4, X7, and X8 to the ground truth transcriptional profiles of the 76 non-neuronal cell-types and neuronal subtypes, we indeed observed a correlation between the deconvolved transcriptional profiles with the ground truth transcriptional profiles of neuronal sub-types (Supplementary Figure S21A). We then sought to evaluate whether increasing the number of deconvolved cell-types could recover the finer neuronal sub-types. We therefore applied STdeconvolve with $K$=76 and were able to identify deconvolved cell-types that were highly correlated in terms of both transcriptional profiles and pixel proportions to finer neuronal subtypes as well as rare cell-types such as pericytes and microglia (Supplementary Figure 21B-C). However, as noted previously, the ability for STdeconvolve to deconvolve neuronal subtypes may not be representative of its expected performance in distinguishing between cell sub-types under less biased gene selection.

### B. Supplementary Methods

#### Deconvolution of simulated MERFISH ST data with STdeconvolve

STdeconvolve was applied to the simulated MERFISH MPOA ST dataset. We selected the model fitted with the lowest perplexity and where the number of "rare" cell-types = 0 resulting in a model with $K$=9 detected cell-types. To compare deconvolved cell-types to the ground truth cell-types in the simulated ST dataset, we computed the Pearson's correlation between every combination of deconvolved cell-type and ground truth cell-type transcriptional profile. Likewise, the Pearson's correlation between the pixel proportions of each deconvolved cell-type and ground truth cell-type was computed. After assignment of deconvolved to ground truth cell-types, the ranking of each gene based on its expression level in the transcriptional profile of the deconvolved or ground truth cell-type for each assigned match was compared.

#### Deconvolution of simulated MERFISH ST data with supervised and semi-supervised reference-based deconvolution approaches

For supervised deconvolution approaches SPOTlight[2] (v0.1.7) and RCTD[4] (v1.2.0), a single cell transcriptomic profile reference was required. To construct this reference, the matrix of gene counts for the 59651 individual cells in the original 12 MPOA tissue sections and their predefined cell-type labels were input into the `seurat` R package[5] (v4.0.1) as recommended in both the SPOTlight and RCTD pipelines. SPOTlight `spotlight_deconvolution` was run using default parameters. RCTD `create.RCTD` was run with CELL_MIN_INSTANCE = 1 and `run.RCTD` was run using default parameters. For supervised method SpatialDWLS[6] (Giotto v2.0.0.953), the simulated ST dataset was normalized and $\log_2$ transformed using default settings. Dimensionality reduction using PCA was performed. Graph-based cluster detection using Louvain clustering[7] was performed using the top 5 principal components with the maximum number of nearest neighbors equal to 100, resulting in the assignment of pixels to 9 clusters. The summed gene expressions of the ground truth cell-types were used as the signature matrix. `runDWLSDeconv` was run using the simulated ST dataset, the transcriptional clusters, and the signature matrix using default settings.

For the semi-supervised approach DSTG[8], we performed the default pipeline to generate pseudo-mixtures of ground truth cells in the scRNA-seq reference dataset, which were used to train the model and deconvolve cell-types in the simulated ST dataset.

To be consistent across approaches, after deconvolution, cell-types in each pixel whose proportions were less than the lowest ground truth pixel proportion for a cell-type (2.5%) were removed, and the remaining cell-type proportions in a pixel were adjusted to sum to 1.

*Deconvolution of simulated MERFISH ST data by reference-based deconvolution approaches using a reference with missing cell-types*

To simulate a single-cell reference with missing cell-types, cells annotated as “excitatory” and “inhibitory” were removed from the previously constructed MERFISH single cell transcriptomic profile single-cell reference and used to train the supervised and semi-supervised methods. The new trained models were then reapplied to the simulated ST MERFISH dataset for deconvolution. Pixel RMSEs were computed based on the deconvolved cell-type proportions and the ground truth dataset, which retained excitatory and inhibitory neuronal cell-types. The same analysis was also repeated for a single-cell reference missing rarer ependymal cells.

*Deconvolution of simulated MERFISH ST data by reference-based deconvolution approaches using a mouse brain scRNA-seq reference*

For the scRNA-seq reference of the mouse cortex, we used the mouse brain scRNA-seq dataset provided by SPOTlight (v0.1.7) containing 1404 cells representing 23 transcriptionally distinct clusters and 34617 genes. This reference was used to train the supervised and semi-supervised approaches.

**Simulating ST data using mixtures of cells from scRNA-seq**

Simulated ST datasets of 900 pixels were generated in which each pixel was the combined gene counts of a mixture of different cell-types up to 8 total individual cells. Individual cells were sampled from cell-types belonging to a scRNA-seq dataset using the original provided annotations[9] (GSE150580).

*Simulating age perturbed ST data*

Individual cells were sampled to generate a simulated ST dataset of "young" luminal cells and macrophages and a simulated ST dataset of "aged" luminal cells and macrophages. Genes detected in more than 5% but less than 100% of pixels were removed. Feature selection was performed to select genes that were significantly overdispersed using a general additive model with a basis of 5 and an adjusted $p$-value cutoff of $1e^{-4}$, which resulted in 877 overdispersed genes for the "young" input corpus. For the "aged" input corpus, overdispersed genes with an adjusted $p$-value cutoff of $1e^{-10}$ were chosen, which resulted in 726 overdispersed genes. STdeconvolve was applied to the "aged" and "young" simulated ST datasets using $K$=2 cell-types and the resulting deconvolved transcriptional profiles were correlated with the average gene expression profiles of the ground truth cell-types of the scRNA-seq dataset using Pearson's correlation. Differentially expressed genes between the annotated deconvolved "young" and "aged" deconvolved macrophage cell-types were determined by dividing the deconvolved transcriptional profiles of the "aged" macrophages by the "young" macrophages and then $\log_2$-transforming the data.

*Deconvolution compared to clustering analysis of simulated cell-type mixtures*

A simulated ST dataset containing pixels with mixtures of 3 cell-types was generated by sampling individual cells from "young" luminal cells, macrophages, and pericytes cell-types of the scRNA-seq reference and combining the gene counts. Genes with less than 10 total reads and detected in less than 10 pixels were removed. For the transcriptional clustering, gene counts were normalized to counts-per-million and $\log_{10}$ transformed with a pseudo-count of 1. Dimensionality reduction using PCA was performed. Graph-based cluster detection using Louvain clustering[7] was performed using the top 30 principal components with the maximum number of nearest neighbors equal to 300, resulting in the assignment of pixels to 2 clusters. For deconvolution with STdeconvolve, we feature selected for genes detected in more than 5% but less than 100% of pixels and were significantly overdispersed using a general additive model with a basis of 5 and an adjusted $p$-value cutoff of 0.05 and used the top 1000 overdispersed genes with the lowest adjusted $p$-values. STdeconvolve was then applied with $K$=3 to deconvolve 3 cell-types. Deconvolved transcriptional profiles were correlated with the average gene expression profiles of the ground truth cell-types of the scRNA-seq dataset using Pearson's correlation and cell-types were matched to ground truth cell-types that had the highest correlations.

*Simulating and deconvolving immune infiltrated vs excluded ST data*

Two simulated ST datasets containing pixels with mixtures of luminal and macrophage cells were generated by sampling individual cells from “young” luminal and “young” macrophage cell-types of the scRNA-seq reference and combining the gene counts. In the “infiltrated” ST simulation, cells were mixed such that the proportion of macrophage followed a gradient from 100% on either the left and right side of the dataset and converged to 0% at the middle. In the “excluded” ST simulation, macrophages were enriched in the 5 columns of pixels on either the left or right and 0% otherwise. For each row, where present, macrophage proportions alternated from 100% to 75%. For both simulations, genes with less than 100 gene counts detected were removed in addition to genes detected in more than 5% but less than 100% of pixels. Feature selection was performed to select genes that were significantly overdispersed using a general additive model with a basis of 5 and an adjusted $p$-value cutoff of 0.05 and the top 800 or 1000 most significant overdispersed genes were kept for either the infiltrated or excluded simulation, respectively. STdeconvolve was then applied to each simulation with $K$=2 to deconvolve 2 cell-types. Deconvolved transcriptional profiles were correlated with the average gene expression profiles of the ground truth cell-types of the scRNA-seq dataset using Pearson’s correlation and cell-types were matched to ground truth cell-types that had the highest correlations.

*Simulating and deconvolving ST data with uniform cell-type distributions*

A simulated ST dataset containing pixels with uniform proportions of 2 cell-types was generated by sampling individual cells from “young” luminal cells and macrophages of the scRNA-seq reference and combining the gene counts. Genes with less than 100 total reads and detected in less than 100 pixels were removed. For deconvolution with STdeconvolve, we feature selected for genes detected in more than 5% but less than 100% of pixels and were significantly overdispersed using a general additive model with a basis of 5 and an adjusted $p$-value cutoff of 0.05 and used the top 1000 overdispersed genes with the lowest adjusted $p$-value. STdeconvolve was then applied with $K$=2 to deconvolve 2 cell-types. Deconvolved transcriptional profiles were correlated with the average gene expression profiles of the ground truth cell-types of the scRNA-seq dataset using Pearson's correlation and cell-types were matched to ground truth cell-types that had the highest correlations.

**Deconvolution of simulated MERFISH mouse brain ST data with BayesSpace and STdeconvolve**

MERFISH data of a coronal section of the mouse brain was obtained[10] (Slice 2 Replicate 1). This dataset contained measured counts of 483 genes for 83546 cells. Transcriptionally distinct cell clusters were identified by normalizing gene counts by counts-per-million, followed by PCA to obtain the top 30 PCs, then graph-based Louvain community detection was applied to obtain 20 clusters. A simulated ST dataset at 100 $\mu m^2$ resolution was generated as described in `*Simulating ST data from single-cell resolution spatially resolved MERFISH data*`, which retained 83247 of the individual cells.

To deconvolve the simulated ST dataset with BayesSpace[11] (v1.3.1), the dataset was first preprocessed setting `platform = "ST"`, using the first 7 principle components (the recommended parameter for ST datasets), setting the number of highly variable genes equal to the number of genes in the dataset (483), and setting `log.normalize` to TRUE. After preprocessing, enhanced spatial clustering into 20 clusters was performed using the first 7 principal components, with

10,000 iterations with the first 100 as burn-in, and default settings otherwise. For an ST dataset, BayesSpace enhanced clustering on ST data divides each pixel into 9 subpixels, each with a cluster membership. We converted the BayesSpace enhanced clustering results into a pixel by "cell-type" proportion matrix by considering each of the 9 subpixels as a cell with a given cell-type (cluster) and computing the proportion of each cell-type in each pixel.

STdeconvolve was applied to the simulated ST dataset setting $K$=20 cell-types. BayesSpace and STdeconvolve predicted pixel proportions were correlated with the ground truth cell-type cluster proportions using Pearson's correlation. Pixel RMSEs were computed based on the deconvolved cell-type proportions and the simulated ST dataset with 20 transcriptional distinct cell-type clusters as ground truth.

**Clustering analysis of ST data of the mouse olfactory bulb (MOB)**

For clustering analysis of the MOB, using the cleaned MOB replicate #8 ST dataset of 260 pixels and 7365 genes, the raw counts were normalized to counts-per-million and adjusted to a $\log_{10}$ scale with pseudo count 1. Subsequently, dimensionality reduction using PCA was performed, and pixels were visualized using 2-D embedding with t-SNE on the top 5 principal components and perplexity = 30. Graph-based cluster detection using Louvain clustering[7] was performed using the top 5 principal components with the maximum number of nearest neighbors equal to 30, resulting in the assignment of pixels to 5 clusters, which were manually annotated based on the physical locations of the pixel clusters on the MOB tissue section[12].

**Deconvolution of ST data of the MOB**

Mouse olfactory bulb datasets were obtained from the original publication[12]. We focused on MOB replicate #8, as the primary MOB ST dataset in this work. We first removed genes with less than 100 reads detected across pixels and pixels with fewer than 100 total gene counts, resulting in a cleaned dataset of 260 pixels and 7365 genes. A gene was overdispersed if the multiple testing adjusted $p$-value was < 0.05 using a general additive model with a basis of 5. For MOB replicate #8, we obtained 255 overdispersed genes. We used STdeconvolve to fit LDA models with a range of integer $K$s from 2 to 20 and chose the model with $K$=12, which was within the range of $K$'s that produced the lowest perplexity and the number of "rare" cell-types with mean pixel proportion < 5% was 0. After deconvolution, cell-types in each pixel whose proportions were less than 5% were removed and the remaining cell-type proportions in each pixel were adjusted to sum to 1.

For supervised approaches SPOTlight, RCTD, and spatialDWLS, we used a previously generated scRNA-seq reference of the MOB[13]. We retained only cells collected from untreated wildtype animals and the resulting matrix encompassed 17709 cells representing 38 previously annotated cell-type clusters and raw counts for 18560 genes. The models were trained as described in `*Deconvolution of simulated MERFISH ST data with supervised and semi-supervised reference-based deconvolution approaches*` and applied to deconvolve cell-types in the cleaned MOB replicate #8 ST dataset of 260 pixels and 7365 genes. After deconvolution, cell-types in each pixel whose proportions were less than 5% were removed and the remaining cell-type proportions were adjusted to sum to 1.

*Comparison of MOB biological replicates*

We applied STdeconvolve to the primary analyzed MOB ST dataset and additional ST datasets of the MOB[12] (replicates 2, 5, and 12, henceforth referred to as MOB-A, -B, and -C, respectively), and compared the transcriptional profiles of deconvolved cell-types between datasets. For each dataset, genes with less than 100 reads across pixels and pixels with less than 100 total reads were discarded. We then selected overdispersed genes for each dataset independent of the other datasets, using a general additive model with a basis of 5 and an adjusted *p*-value cutoff of 0.05. This resulted in input datasets of 279 pixels by 385 genes MOB-A), 267 pixels by 162 genes (MOB-B), 260 pixels by 255 genes (primary MOB), and 278 pixels by 177 genes (MOB-C). When fitting LDA models to each dataset using STdeconvolve, $K$=12 was chosen for each dataset because the number of cell-types with low proportions across the pixels was consistently stable until this threshold for all datasets.

*Deconvolution of ST data of the MOB by reference-based deconvolution approaches using a reference with missing cell-types*

To simulate a single-cell reference with missing cell-types, cells of the MOB scRNA-seq reference that were part of "OEC" clusters 1-5 were removed. Supervised approaches were trained using this new reference and the newly trained models were then reapplied to the cleaned MOB replicate #8 ST dataset of 260 pixels and 7365 genes for deconvolution. After deconvolution, cell-types in each pixel whose proportions were less than 5% were removed and the remaining cell-type proportions were adjusted to sum to 1.

*Identification of MOB cell-type putative marker genes*

`Seurat::FindAllMarkers()` with default parameters was applied to the raw counts of the MOB scRNA-seq reference to obtain differentially expressed genes for each cell-type cluster relative to all other cell-type clusters. Putative cell-type marker genes were defined as those with adjusted $p$-values < 0.05 and average $\log_2$ fold-change > 1. Putative cell-type marker genes were ranked by lowest adjusted $p$-value followed by largest average $\log_2$ fold-change. For OEC putative markers, the same procedure was followed except that cells in OEC cell-type clusters 1 through 5 were combined into a single OEC cluster.

*Deconvolution of ST data of the MOB by reference-based deconvolution approaches using a mouse brain scRNA-seq reference*

For the scRNA-seq reference of the mouse brain, we used the scRNA-seq dataset provided by SPOTlight (v0.1.7) containing 1404 cells representing 23 transcriptionally distinct clusters and 34617 genes. Supervised and semi-supervised methods were trained using this reference and the newly trained models were then reapplied to the cleaned MOB replicate #8 ST dataset of 260 pixels and 7365 genes for deconvolution. After deconvolution, cell-types in each pixel whose proportions were less than 5% were removed and the remaining cell-type proportions were adjusted to sum to 1.

*Comparing deconvolution of the MOB between STdeconvolve, SPOTlight, RCTD, and spatialDWLS*

Deconvolved transcriptional profiles were not returned by the supervised methods, we compared methods to STdeconvolve by evaluating the Pearson's correlation between the pixel proportions of STdeconvolve deconvolved cell-types and cell-types from another method. Predicted cell-types of the MOB reference that were deconvolved by a supervised method were matched to STdeconvolve cell-types that had the highest Pearson's correlation.

*Identification of VLMC cell-type putative marker genes*

The procedure described in `*Identification of MOB cell-type putative marker genes*` was followed using the mouse brain scRNA-seq dataset described in `*Deconvolution of ST data of the MOB by reference-based deconvolution approaches using a mouse brain scRNA-seq reference*`.

**Deconvolution of DBiT-seq data with STdeconvolve**

DBiT-seq dataset of an E11 mouse embryo lower body sample (GSM4364242_E11-1L) was obtained from the original publication[14]. Pixels with less than 100 gene counts, genes with less than 100 total counts, and mitochondrial genes were removed resulting in a filtered dataset of 1831 pixels and 7171 genes. We then feature selected for the top 1000 most significant overdispersed genes using a general additive model with a basis of 5 and an adjusted *p*-value cutoff of 0.01 that were detected in more than 5% but less than 100% of pixels. STdeconvolve was applied to the final ST dataset with $K$=13 cell-types, equal to the number of transcriptional clusters for this sample in the original publication. After deconvolution, cell-types in each pixel whose proportions were less than 5% were removed and the remaining cell-type proportions were adjusted to sum to 1.

**Deconvolution of Slide-seq data**

A Slide-seq puck (Puck_180819_12) of the mouse cerebellum was obtained from the original publication[15].

*With STdeconvolve*

Pixels with less than 50 gene counts, and genes with less than 100 total counts that were detected in less than 90% of pixels were removed. We feature selected for the top 400 significantly overdispersed genes using a general additive model with a basis of 5 and an adjusted *p*-value cutoff of 0.01. The final input dataset for STdeconvolve was 23490 beads and 400 genes. We used STdeconvolve to fit LDA models with a range of integer *K*s from 2 to 20 and chose the model with *K*=14, which was the highest *K* within the range of *K*'s that produced the lowest perplexity and the number of "rare" cell-types with mean pixel proportion < 5% was 0. Each deconvolved cell-type was annotated by matching its transcriptional profile with a cell-type subcluster in a previously annotated DropSeq scRNA-seq reference of the mouse cerebellum[16] (BrainCellAtlas version 2018.04.01). This was done by generating transcriptional profiles of the scRNA-seq reference cell-types by grouping cells into each subcluster and combining the genes counts. Next, the Pearson's correlation between every combination of deconvolved and ground truth cell-type transcriptional profile was computed and deconvolved cell-types were annotated as the ground truth cell-type that had the highest Pearson's correlation between their transcriptional profiles.

*With RCTD*

RCTD was applied to the full original puck of 19782 genes and 46376 beads with the mouse cerebellum DropSeq scRNA-seq dataset described above as the reference. RCTD `create.RCTD` was run with CELL_MIN_INSTANCE = 1 and `run.RCTD` was run using default parameters in doublet mode, which resulted in the deconvolution of 11330 of the beads, of which 11320 were also deconvolved by STdeconvolve.

*Comparison of STdeconvolve and RCTD*

3583 Slide-seq beads identified as singlets by RCTD and were also deconvolved by STdeconvolve were kept. For RCTD, the first cell-type was considered the cell-type deconvolved in the bead. For STdeconvolve, the cell-type with the largest predicted proportion was considered the cell-type deconvolved in the bead. The overlaps of beads determined to contain Purkinje neurons or Bergmann glia by either RCTD or STdeconvolve were determined and Fisher's exact test was performed to calculate statistically significant enrichment.

**Deconvolution of 10X Visium data with STdeconvolve**

A 10X Visium dataset of a coronal section of the mouse cortex was obtained[17]. Pixels with less than 100 gene counts and genes with less than 100 total counts were removed resulting in a filtered dataset of 2702 pixels and 13548 genes. We then feature selected for the top 1000 most significant overdispersed genes using a general additive model with a basis of 5 and an adjusted *p*-value cutoff of 0.01 and were detected in more than 5% but less than 100% of pixels. STdeconvolve was applied to the final ST dataset with $K$=20 cell-types.

**Clustering analysis of breast cancer sections**

For clustering analysis of the breast cancer sections, we took the combined dataset of 1029 pixels and 372 overdispersed genes using a general additive model with a basis of 5 and an adjusted *p*-value cutoff of 0.05, and $\log_{10}$ transformed with pseudo count of 1, and dimensionality reduction using PCA was performed. In a manner similar to the original publication, pixels were clustered into 3 groups using the "Ward.D" method and the Euclidean distance calculated using the first 2 principle components This resulted in the assignment of pixels to 3 clusters which corresponded to the annotations of the 3 histological sections previously annotated by pathologists[18].

**Deconvolution of ST data of breast cancer sections with STdeconvolve**

ST datasets of 4 breast cancer sections were obtained from the original publication[18]. Genes with less than 10 reads across pixels or pixels with less than 10 total reads were removed from each dataset and genes present in more than 95% of pixels for given dataset were removed. Overdispersed genes were determined for each dataset as described in ` *Deconvolution of ST data of the MOB* ` using the same parameters. After, we combined the 4 breast cancer datasets into a single dataset of 1029 pixels with counts for 372 genes found to be overdispersed in at least one dataset. We trained LDA models on this combined dataset with STdeconvolve using a range of $K$ from 2 to 20 and selected $K$=15.

*<u>Gene set enrichment analysis of deconvolved breast cancer cell-types</u>*

To interpret the transcriptional profiles of the deconvolved cell-types in ST data of breast cancer sections, we used gene set enrichment analysis as implemented in the `liger` R package[19]. We filtered the list of 16771 Homo sapiens Gene Ontology gene set terms[20] to include those which contained at least 1 gene present in the input ST dataset corpus used with STdeconvolve, resulting in 4238 terms. We then performed iterative gene set enrichment analysis on the ranked expression profile of the genes as previously described in `*Annotation and matching of deconvolved and ground truth cell-types*`

*Enrichment of deconvolved cell-types in breast cancer pathological labels*

A one-sided Fisher's exact test was performed to test if a deconvolved cell-type was significantly enriched in pixels that also labeled with a given pathological annotation. A cell-type was said to be in a pixel if its proportional contribution was > 0.1.

**Accuracy of STdeconvolve with respect to ST dataset size**

We assessed the performance of STdeconvolve using the simulated MERFISH MPOA ST dataset and measured RMSE with respect to varying the number of pixels. We randomly sampled pixels from the simulated ST dataset up to a given number of pixels and used this sub-sampled dataset as input into STdeconvolve, with $K$=9. The mean RMSE of the deconvolved sub-sampled pixels was determined with respect to the ground truth cell-type proportions of the sampled pixels. We repeated this process 10 times for pixel subsample sizes of 5, 10, 50, 100, and 150.

### C. Supplementary Figures

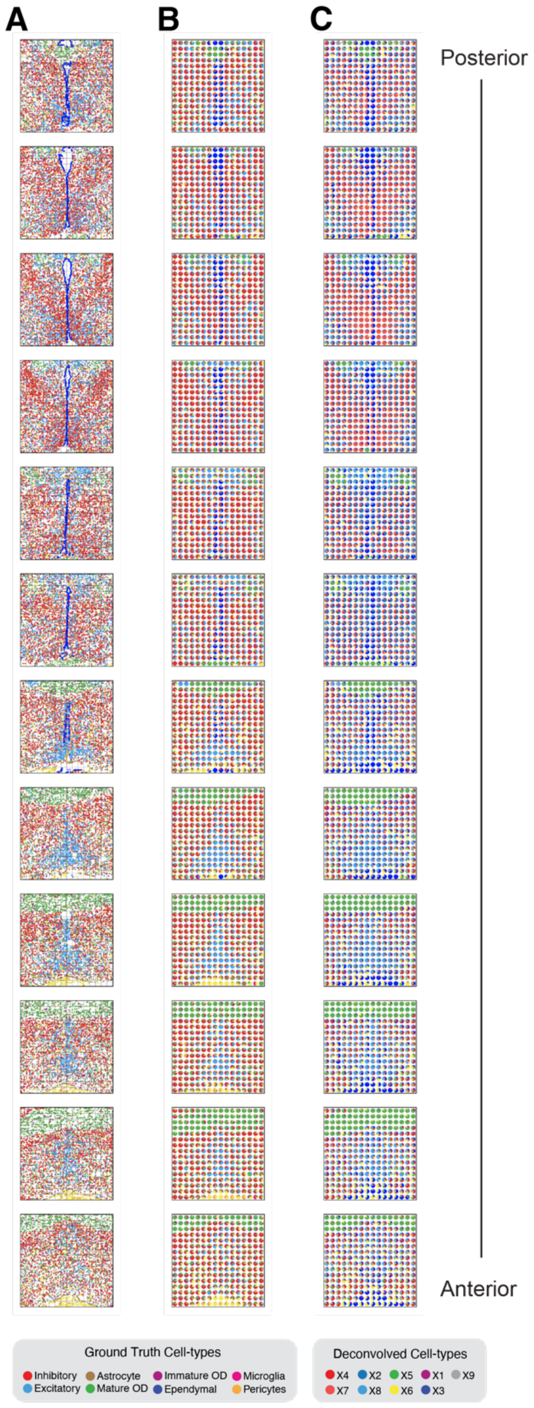


**Figure S1. Simulated MERFISH MPOA ST data and STdeconvolve results for 9 deconvolved cell-types across all 12 tissue sections.** A) Spatial positions of individual cells of each 9 major cell-types across tissue sections. Simulated pixels demarcated as dashed line squares. B) Simulated pixels as pie charts indicating the ground truth pixel proportions of each 9 major cell-types across tissue sections. C) STdeconvolve pixel proportion predictions of 9 deconvolved cell-types.

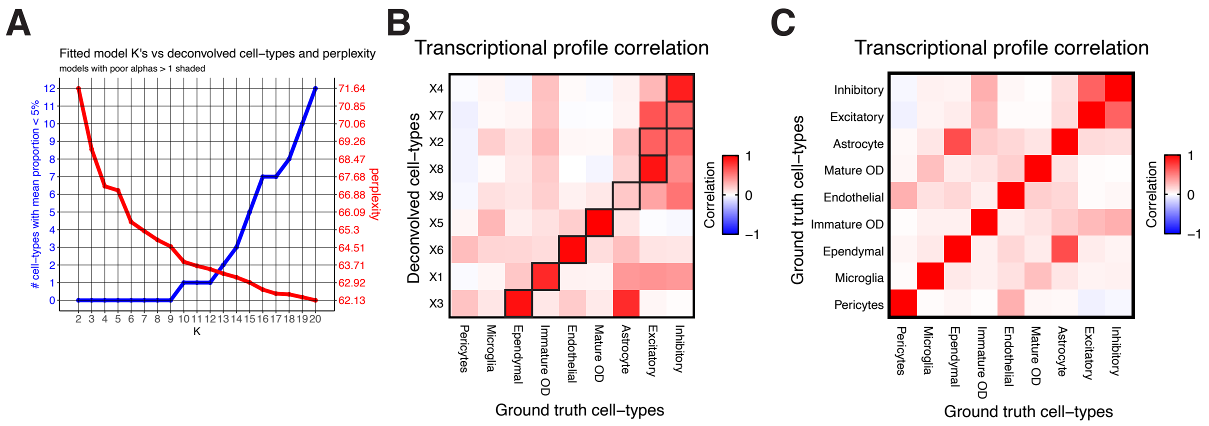


**Figure S2. Comparison of deconvolved cell-types from STdeconvolve to ground truth cell-types of simulated MERFISH MPOA ST data.** A) STdeconvolve *K* versus "rare" cell-types and perplexity. B) Pearson's correlation between the transcriptional profiles of the 9 ground truth cell-types in the MERFISH MPOA data and the 9 deconvolved cell-types by STdeconvolve. C) Pearson's correlation between the transcriptional profiles of the 9 ground truth cell-types.

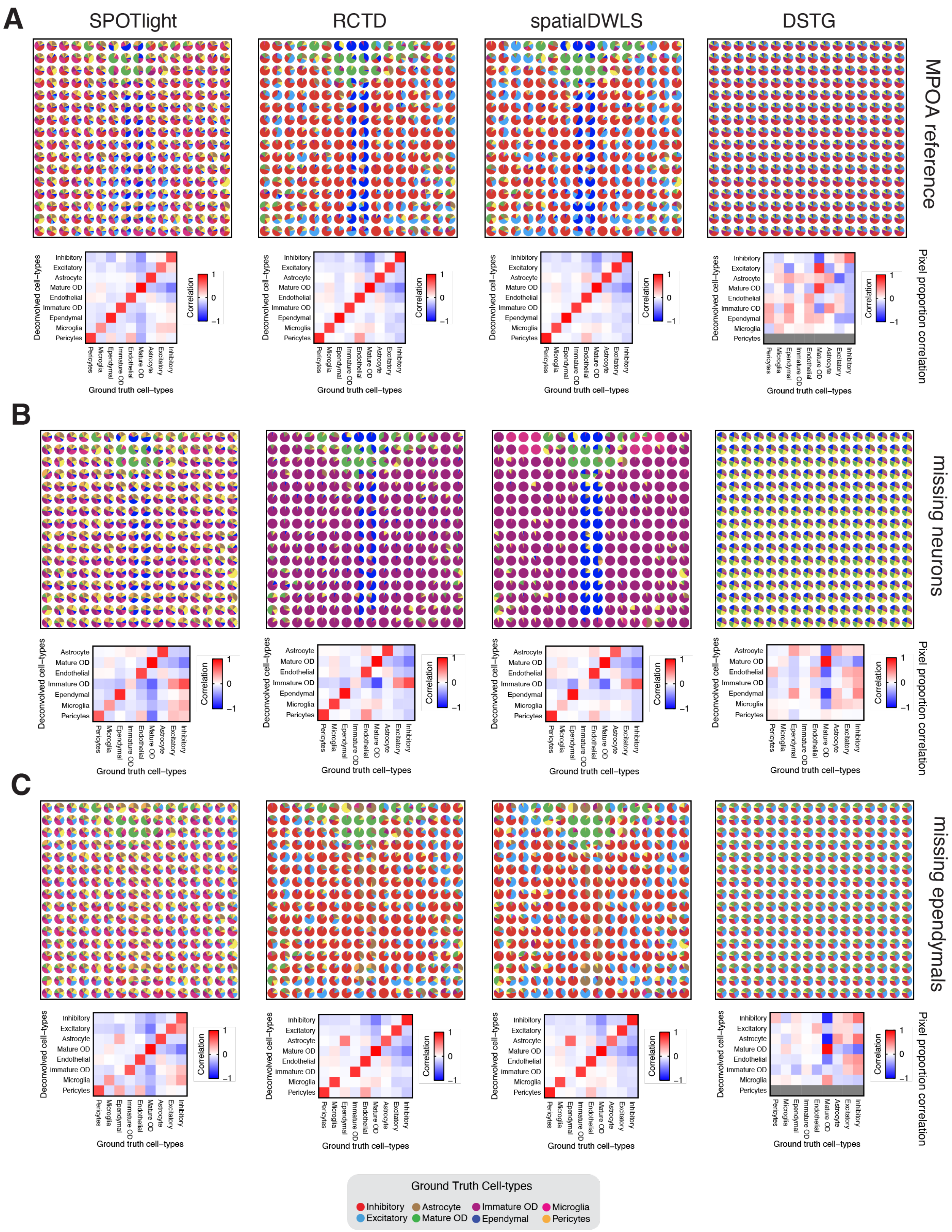
A
SPOTlight
RCTD
spatialDWLS
DSTG
MPOA reference
Pixel proportion correlation
B
missing neurons
Pixel proportion correlation
C
missing ependymals
Pixel proportion correlation
Deconvolved cell−types
Ground truth cell−types
Correlation
Inhibitory
Excitatory
Astrocyte
Mature OD
Endothelial
Immature OD
Ependymal
Microglia
Pericytes
Ground Truth Cell-types
Inhibitory
Excitatory
Astrocyte
Mature OD
Immature OD
Ependymal
Microglia
Pericytes

**Figure S3. Cell-type predictions of supervised and semi-supervised deconvolution approaches across simulated MERFISH MPOA ST dataset pixels with different single-cell transcriptomics references.** A) Deconvolved pixel proportions of cell-types by each supervised and semi-supervised deconvolution approach using the MERFISH MPOA single-cell transcriptomics data as the reference. B) Deconvolved pixel proportions of cell-types by each supervised and semi-supervised deconvolution approach using the MERFISH MPOA single-cell transcriptomics data with missing neurons as the reference. C) Deconvolved pixel proportions of cell-types by each supervised and semi-supervised deconvolution approach using the MERFISH MPOA single-cell transcriptomics data with missing ependymal cells as the reference.

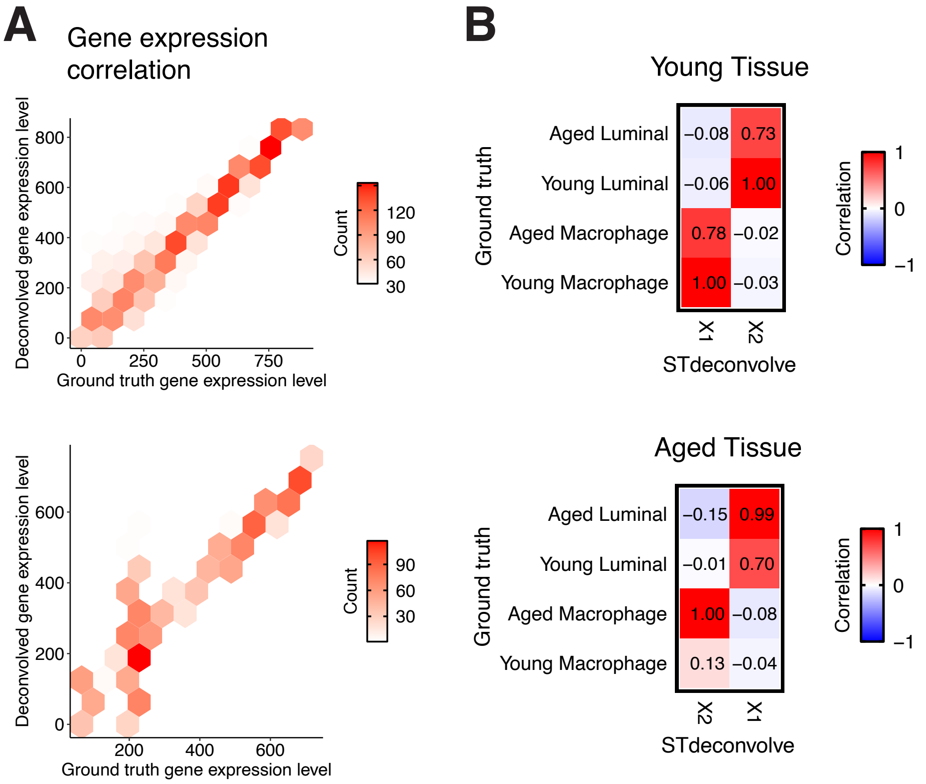


**Figure S4. STdeconvolve deconvolves simulated ST data of aged and young tissues.** A) The ranking of each gene based on its expression level in the transcriptional profiles of the deconvolved cell-types, compared to its gene rank in the transcriptional profile of the matched ground truth cell-type for simulated ST data of young (top) and aged (bottom) tissues. B) Heatmap of Pearson's correlations between the transcriptional profiles of the ground truth cell-types and deconvolved cell-types from the simulated ST data of young (top) and aged (bottom) tissues.

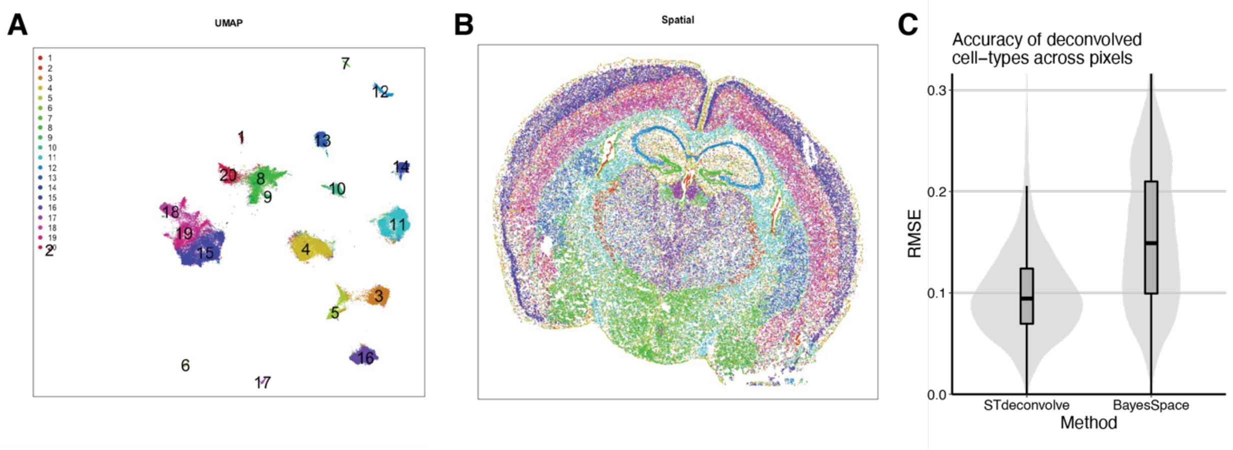


**Figure S5. Single-cell transcriptional clustering of MERFISH data of a coronal section of the mouse brain.** A) UMAP embedding of single cells colored by transcriptional cluster assignment. B) Spatial positions of cells colored by transcriptional cluster assignment. C) Root-mean-square-error (RMSE) of the deconvolved cell-type proportions compared to single-cell clustering for the MERFISH mouse brain data.

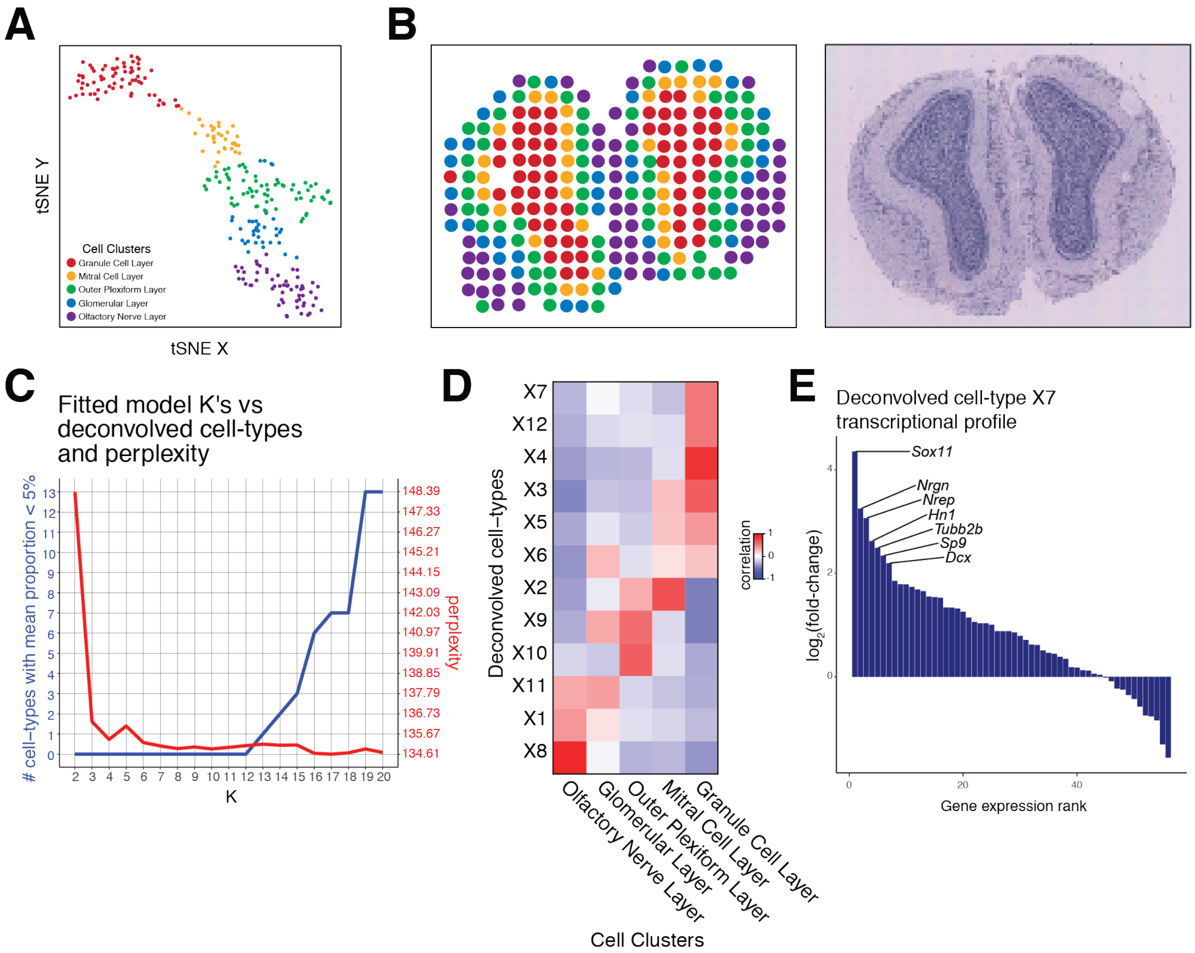


**Figure S6. Additional of the MOB.** A) tSNE plot of the MOB pixels colored by their transcriptional cluster memberships. B) Visualization of MOB pixels colored by their transcriptional cluster memberships on the pixel spatial coordinates (left). H&E-stained image of the corresponding MOB tissue section (right). C) STdeconvolve *K* versus "rare" cell-types and perplexity. D) Pearson's correlation between the pixel proportions of 12 deconvolved cell-types by STdeconvolve and the MOB pixel transcriptional cluster memberships. E) $Log_2$ fold-change of the deconvolved transcriptional profile genes of deconvolved cell-type X7 with respect to the mean deconvolved expression of the other 11 deconvolved cell-types.

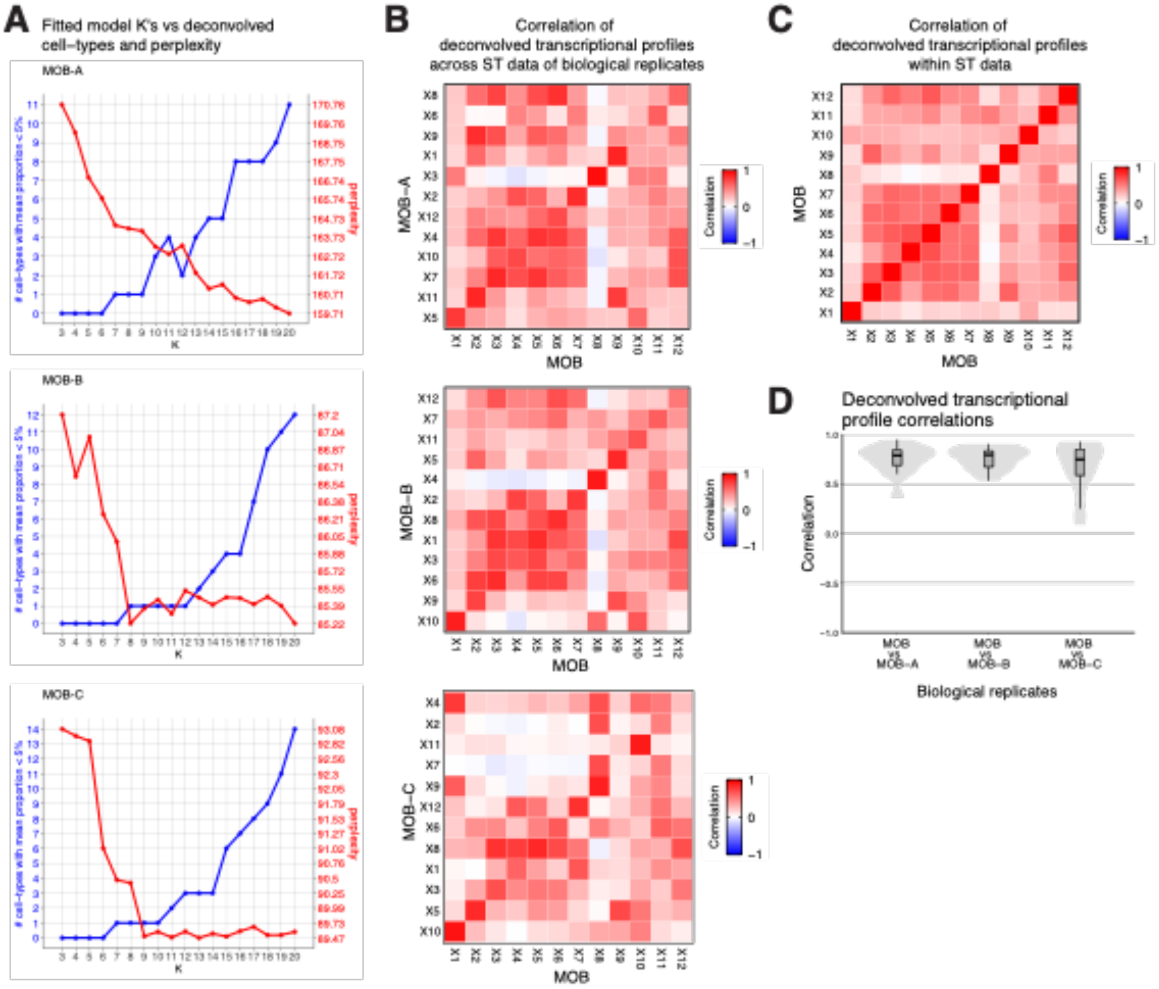


**Figure S7. Stability of deconvolved cell-types across MOB ST data of biological replicates.** A) STdeconvolve *K* versus “rare” cell-types and perplexity for MOB-A, MOB-B, and MOB-C. B) Pearson’s correlations between deconvolved transcriptional profiles of deconvolved cell-types of the primary MOB ST data versus MOB-A, MOB-B, and MOB-C ST data. C) Pearson’s correlations between deconvolved transcriptional profiles of deconvolved cell-types within the primary MOB ST data. D) Distribution of Pearson’s correlations between deconvolved transcriptional profiles of corresponding matched deconvolved cell-types between the primary MOB ST data versus MOB-A, MOB-B, and MOB-C ST data.

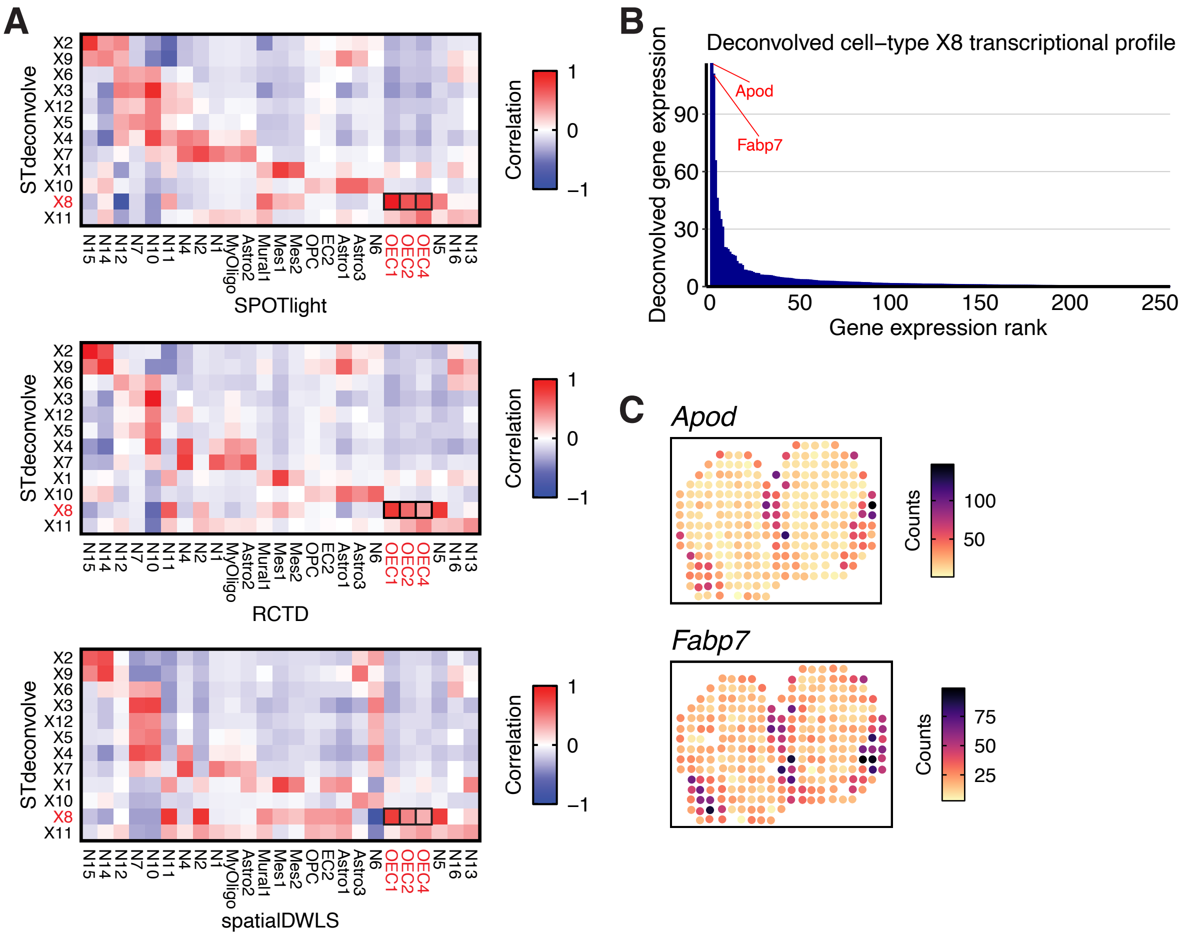


**Figure S8. Correlations between deconvolved cell-types of the MOB ST dataset by STdeconvolve and supervised deconvolution approaches.** A) Heatmaps of the Pearson's correlations between pixel proportions of deconvolved cell-types of different methods. Black squares highlight correlations between STdeconvolve cell-type X8 and OEC cell-types predicted by supervised approaches. B) Transcriptional profile of STdeconvolve cell-type X8. Top expressed genes include *Apod* and *Fabp7*, which were also top differentially expressed putative marker genes of the OEC cell cluster. C) Gene counts in each pixel of the MOB ST dataset of top putative marker genes for OEC cell cluster.

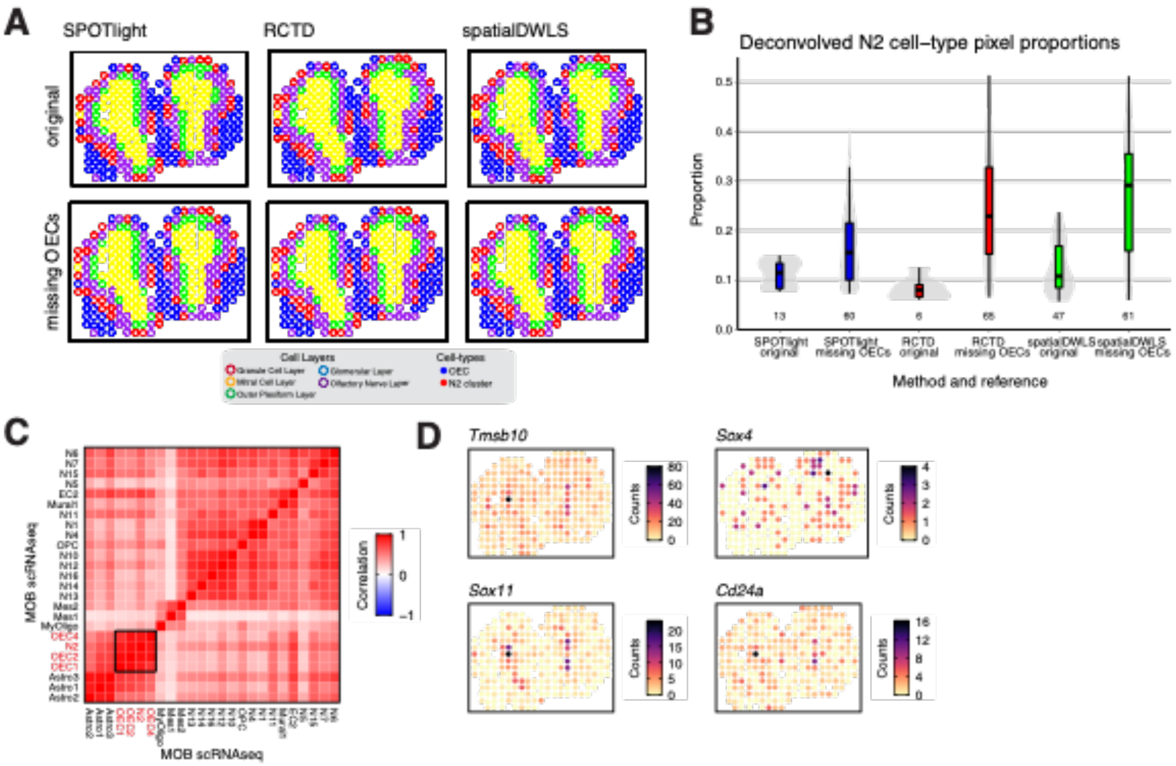


**Figure S9. Effect of missing OEC cell cluster in MOB reference on supervised deconvolution approaches.** A) Pixels of the MOB ST dataset represented as pie charts indicating the deconvolved pixel proportions of OEC clusters (blue) and the N2 cell cluster (red), by supervised deconvolution approaches, trained with either the full MOB scRNA-seq reference or the reference missing OEC clusters. B) Deconvolved pixel proportions of N2 cell cluster by either SPOTlight (blue), RCTD (red), or spatialDWLS (green) trained with either the full MOB scRNA-seq reference or the reference missing OEC clusters. Numbers below each boxplot indicate the number of pixels in the MOB ST dataset for which N2 cluster was predicted. C) Pearson’s correlations between transcriptional profiles of cell-types in the MOB scRNA-seq reference. D) Gene counts in each pixel of the MOB ST dataset of top putative marker genes for N2 cell cluster.

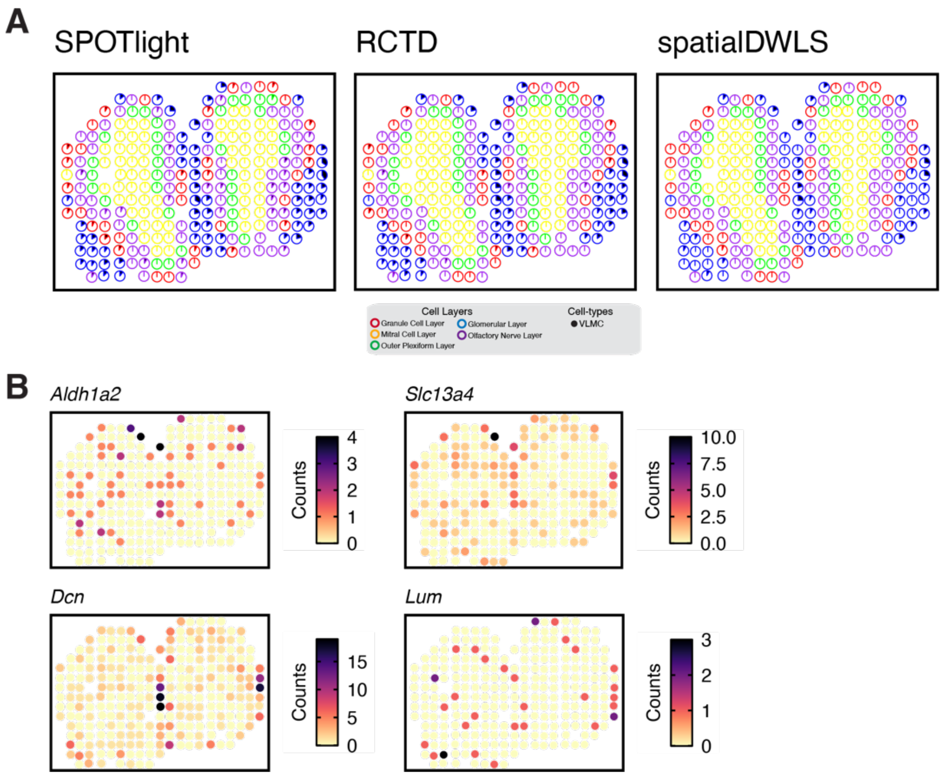


**Figure S10. Misassignment of VLMC cluster in MOB ST data by supervised deconvolution approaches when trained with cortex scRNA-seq reference.** A) Pixels of the MOB ST dataset represented as pie charts indicating the deconvolved pixel proportions of VLMC cluster (black) by either SPOTlight, RCTD, or spatialDWLS. B) Genes counts of putative top marker genes of VLCM based on the scRNA-seq reference and *Lum*, a unique marker of VLMCs[21], in each pixel of the MOB ST dataset.

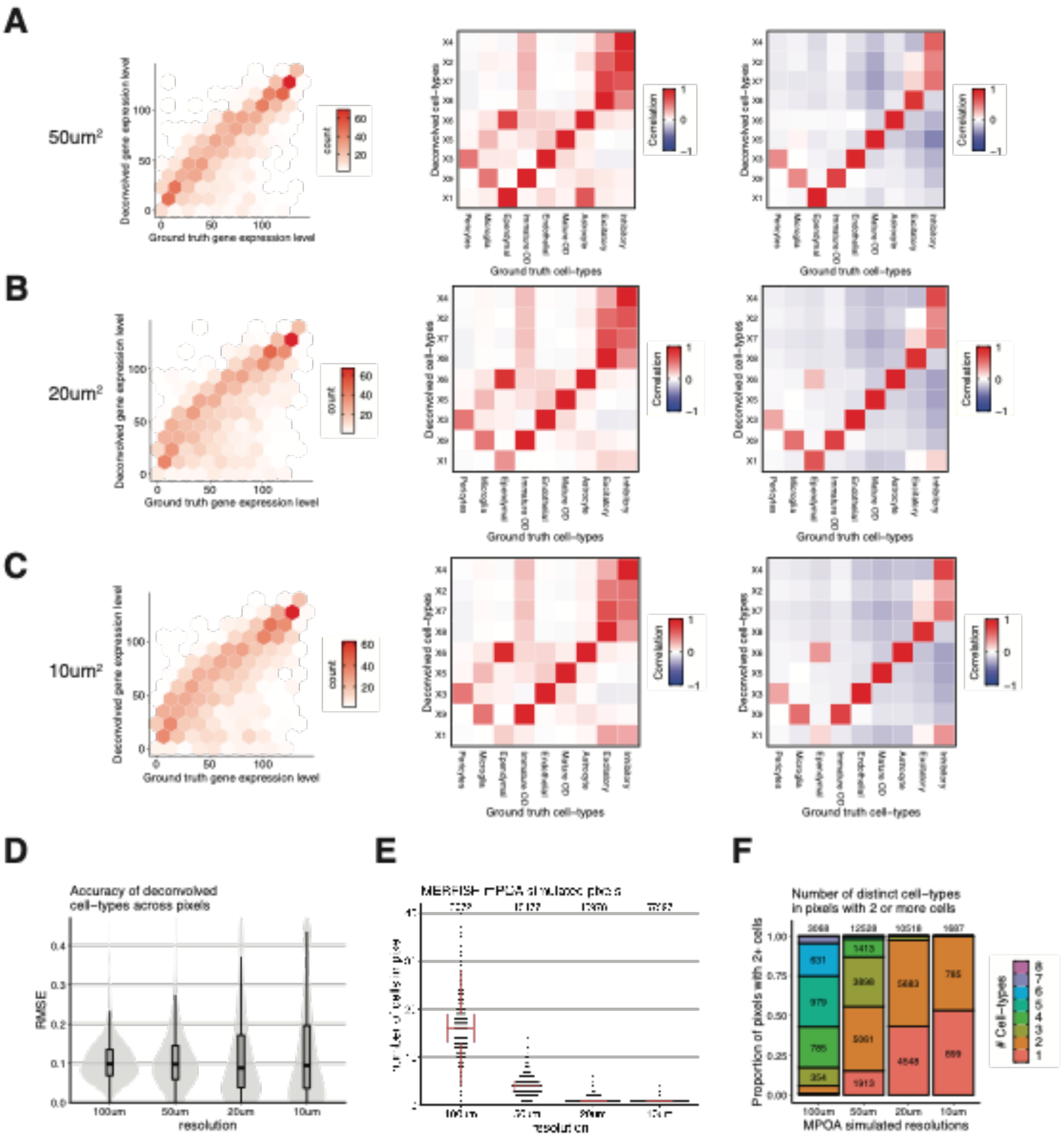


**Figure S11. Comparison of STdeconvolve cell-types to ground truth cell-types in high resolution pixel MERFISH MPOA ST datasets.** A) Deconvolution accuracy in 50 µm$^2$ simulated pixels. Left: The ranking of each gene based on its expression level in the transcriptional profiles of the deconvolved cell-types, compared to its gene rank in the transcriptional profile of the matched ground truth cell-type. Middle: Pearson's correlation between the transcriptional profiles of the 9 ground truth cell-types in MERFISH MPOA data and 9 deconvolved cell-types by STdeconvolve. Right: Pearson's correlation between the pixel proportions of the 9 ground truth cell-types in the MERFISH MPOA data and 9 deconvolved cell-types by STdeconvolve. B) Deconvolution accuracy in 20 $\mu m^2$ simulated pixels. C) Deconvolution accuracy in 10 $\mu m^2$ simulated pixels. D) Root-mean-square-error (RMSE) of the deconvolved cell-type pixels proportions compared to ground truth cell-types at each resolution of the simulated ST dataset. E) Distributions of number of individual cells contained within pixels of each simulated resolution of the MERFISH MPOA ST dataset. Numbers above each dataset indicate the total number of pixels for each simulated ST data. F) Stacked bar plots showing the fraction of pixels that contain a distinct number of different cell-types for each simulated resolution of the MERFISH MPOA ST dataset. This was done for pixels that contain at 2 or more individual cells. Numbers within each box indicate the total number of pixels with that many distinct cell-types and number above the bars indicate the total number of pixels with 2 or more individual cells.

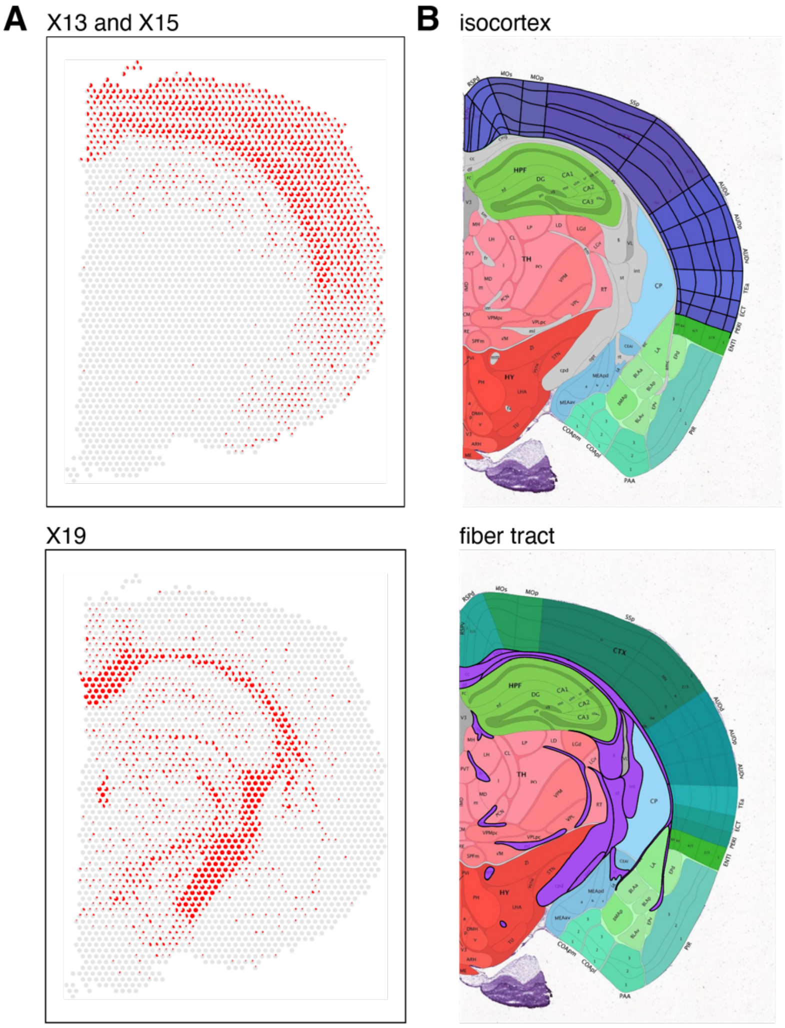


**Figure S12. STdeconvolve pixel proportion of select deconvolved cell-types in 10X Visium data of the mouse brain.** A) STdeconvolve pixel proportions for select deconvolved cell-types. B) Visually corresponding annotated brain region from the Allen Brain Atlas[22].

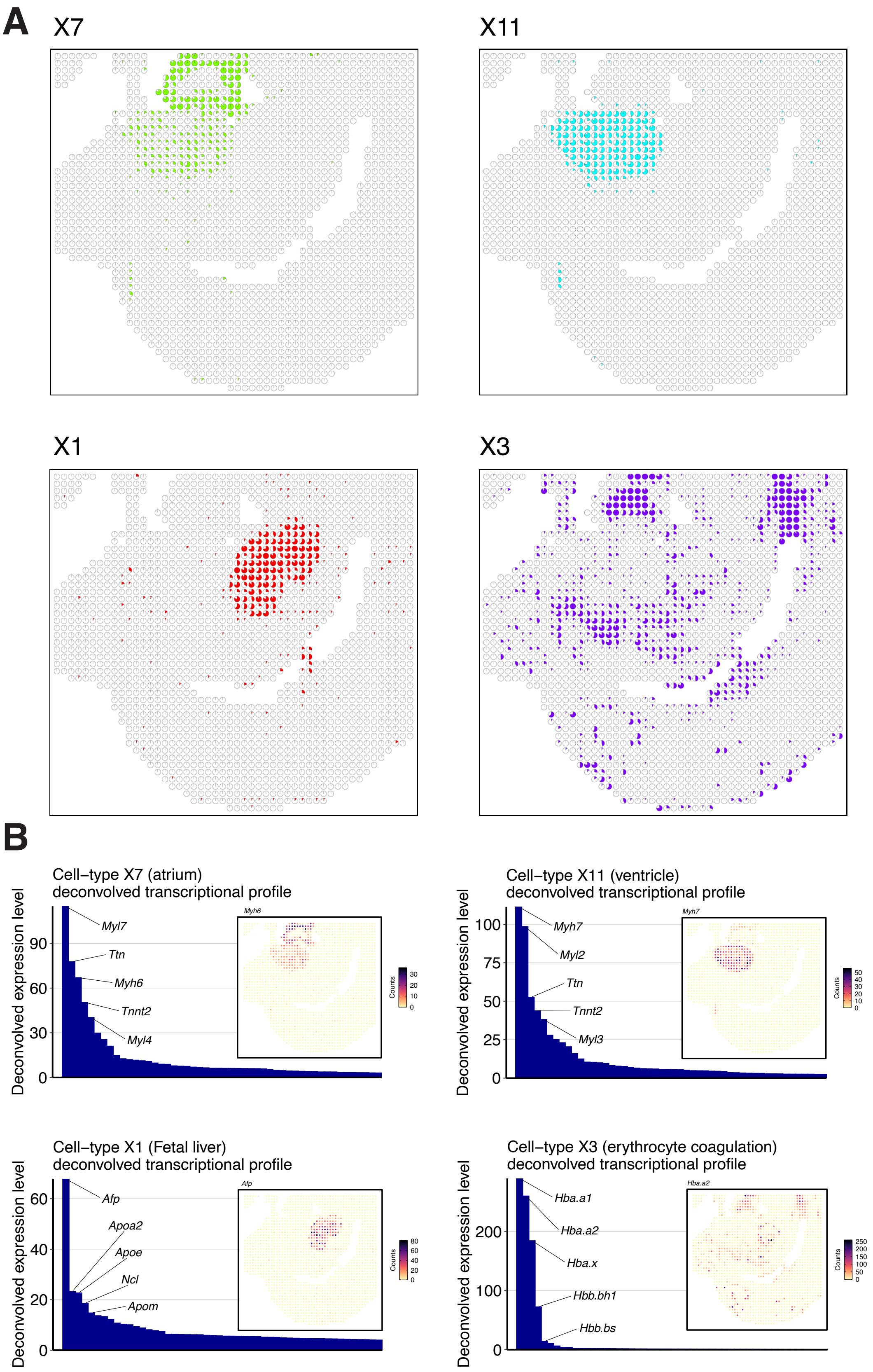
A
X7
X11
X1
X3
B
Cell−type X7 (atrium)
deconvolved transcriptional profile
Deconvolved expression level
Myl7
Ttn
Myh6
Tnnt2
Myl4
Cell−type X11 (ventricle)
deconvolved transcriptional profile
Myh7
Myl2
Ttn
Tnnt2
Myl3
Cell−type X1 (Fetal liver)
deconvolved transcriptional profile
Afp
Apoa2
Apoe
Ncl
Apom
Cell−type X3 (erythrocyte coagulation)
deconvolved transcriptional profile
Hba.a1
Hba.a2
Hba.x
Hbb.bh1
Hbb.bs
Counts

**Figure S13. STdeconvolve pixel proportions and transcriptional profiles of select deconvolved cell-types in DBiT-seq data of an E11 mouse embryo lower tail section.** A) Pixel proportions of select deconvolved cell-types X7, X11, X1, and X3 corresponding to the atrium, ventricle, fetal liver, and erythrocyte coagulation, respectively, annotated in the previous publication[14]. B) Deconvolved transcriptional profiles for cell-types X7, X11, X1, and X3 with the expression of the top gene in each deconvolved transcriptional profile visualized in the original tissue (inset).

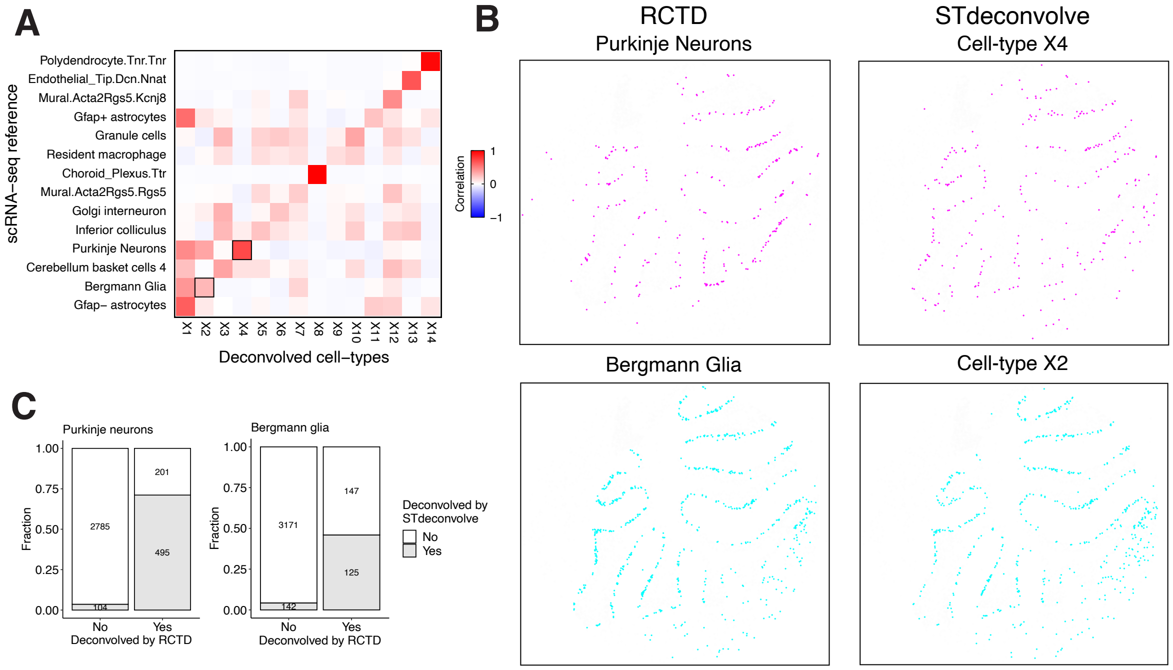


**Figure S14. STdeconvolve predicted cell-type proportions of Purkinje neurons and Bergmann glia in Slide-seq data of the mouse cerebellum compared to RCTD.** A) Pearson's correlation between the deconvolved transcriptional profiles of the 14 STdeconvolve cell-types and best matched cell-types in the mouse cerebellum scRNA-seq reference[16]. B) Beads predicted to contain Purkinje neurons (cyan) or Bergmann glia (magenta) by either RCTD (left) or STdeconvolve (right). C) Stacked bar plots indicating the fractions of beads predicted to contain Purkinje neurons (left) or Bergmann glia (right) by RCTD and STdeconvolve.

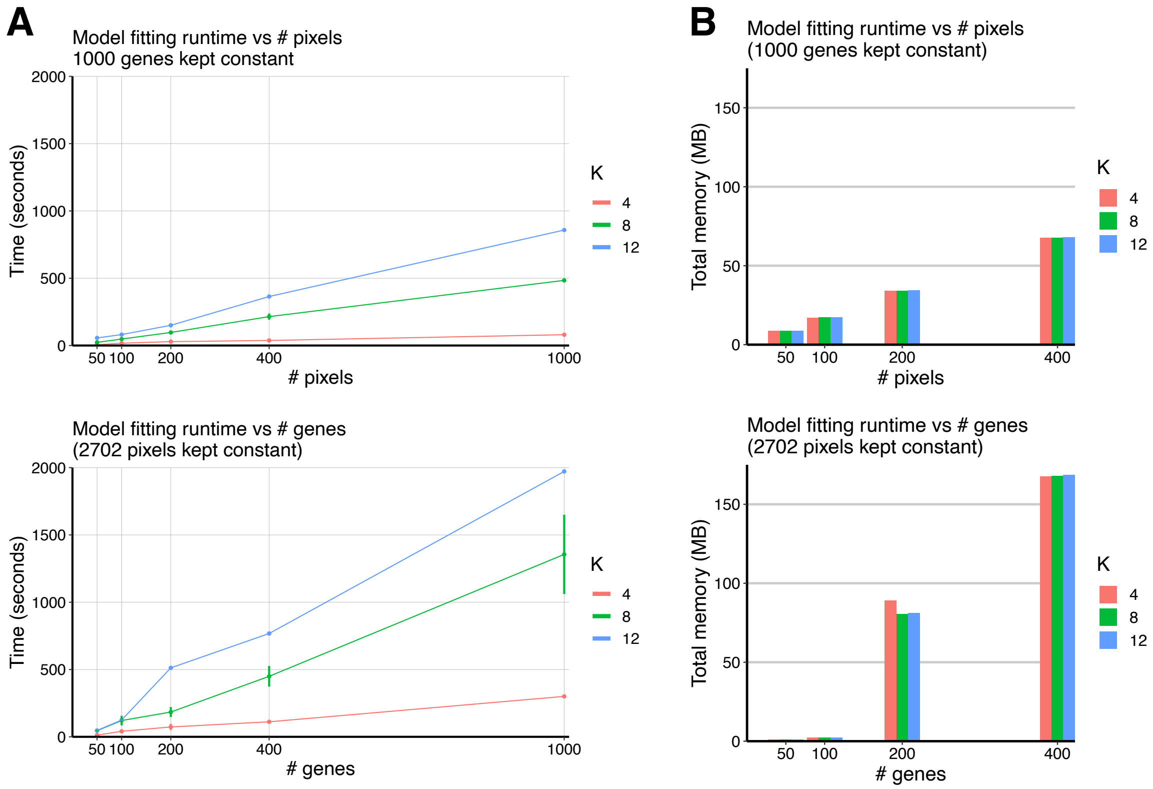


**Figure S15. Runtime and memory usage by STdeconvolve.** A) Runtime as a function of dataset size in terms of the number of pixels (top) or genes (bottom), and number of cell-types *K*. For scaling pixels, the top 1000 most significant overdispersed genes were selected and kept constant. For scaling genes, all 2702 pixels in the dataset were used and kept constant. B) Memory usage as a function of dataset size in terms of scaling pixels (top) or genes (bottom), and number of cell-types *K*. Again, for scaling pixels, the top 1000 most significant overdispersed genes were selected and kept constant. For scaling genes, all 2702 pixels in the dataset were used and kept constant.

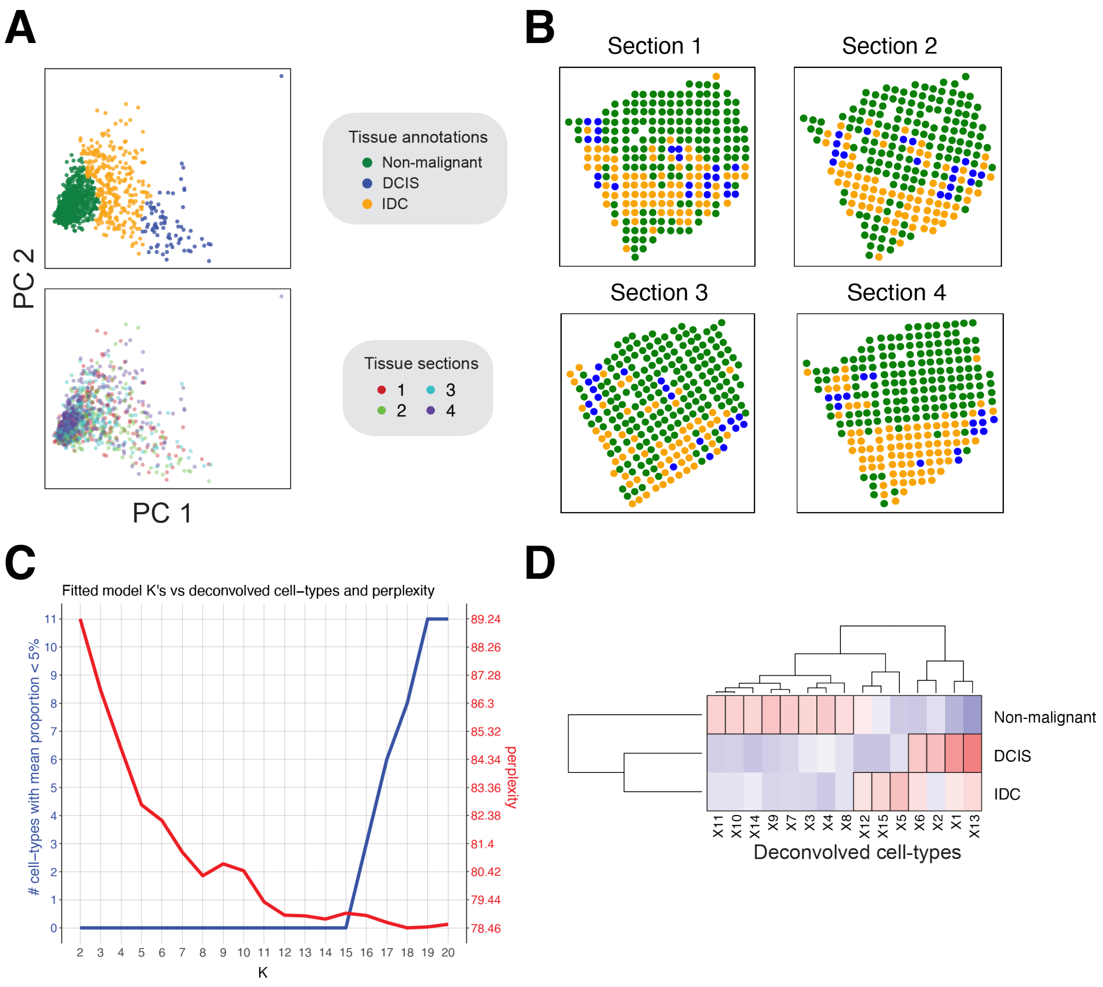


**Figure S16. Deconvolution of the breast cancer ST data by STdeconvolve.** A) Transcriptional clustering of ST pixels. PCA plot of the 1029 breast cancer pixels of all 4 sections. Pixels colored by cluster corresponding to a tissue annotation (top) or to the associated tissue section (bottom). B) Pixels of each breast cancer tissue sections colored by tissue annotation. C) STdeconvolve *K* versus "rare" cell-types and perplexity. D) Pearson's correlation between the pixel proportions of 15 deconvolved cell-types by STdeconvolve and the breast cancer tissue annotations. Black boxes highlight tissue annotations with highest correlations to each deconvolved cell-type.

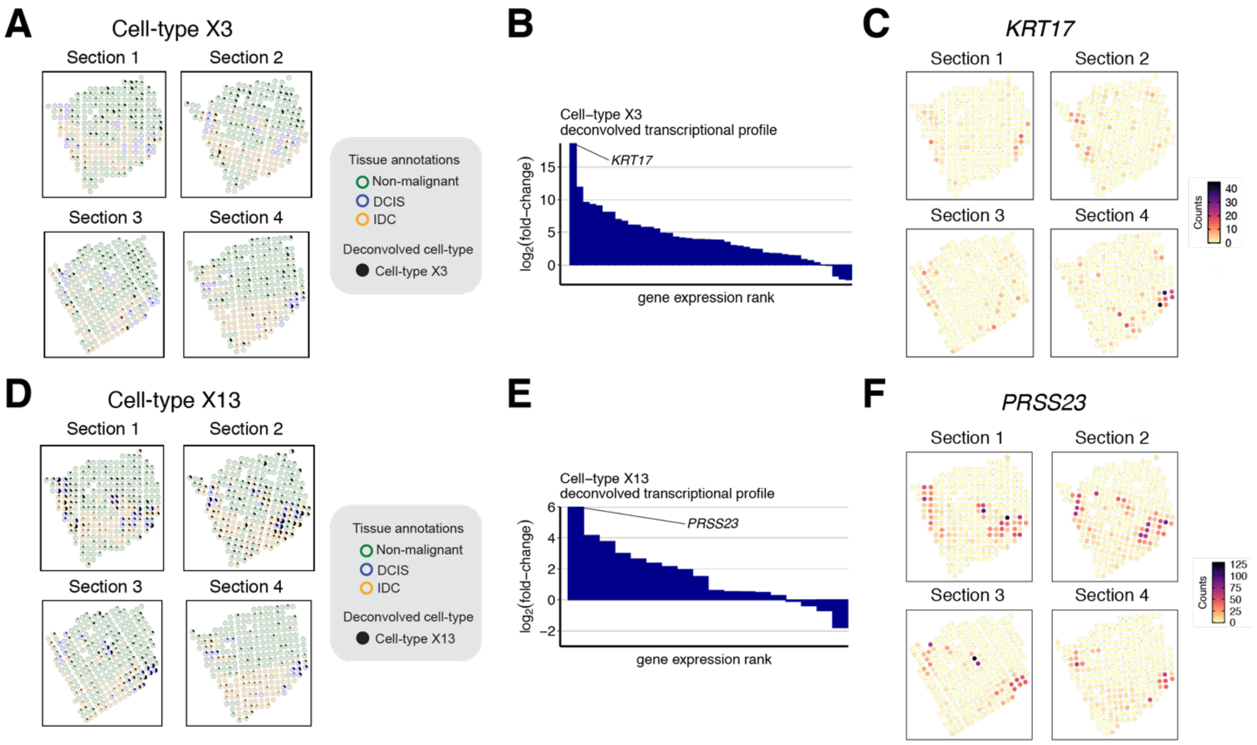


**Figure S17. STdeconvolve cell-types X3 and X13 of the breast cancer ST data.** A) Pixels of each breast cancer tissue section with outlines colored corresponding to their assigned tissue annotation. Pixels represent pie charts of cell-type pixel proportions, where the pixel proportion of deconvolved cell-type X3 is colored in black. B) Log$_2$ fold-change of the deconvolved transcriptional profile for deconvolved cell-type X3 with respect to the mean deconvolved transcriptional profile for the other 14 deconvolved cell-types. C) Gene counts of *KRT17*, the top differentially upregulated gene for cell-type X3, in each pixel of the breast cancer tissue sections. D-F) Same as (A-C) but for cell-type X13.

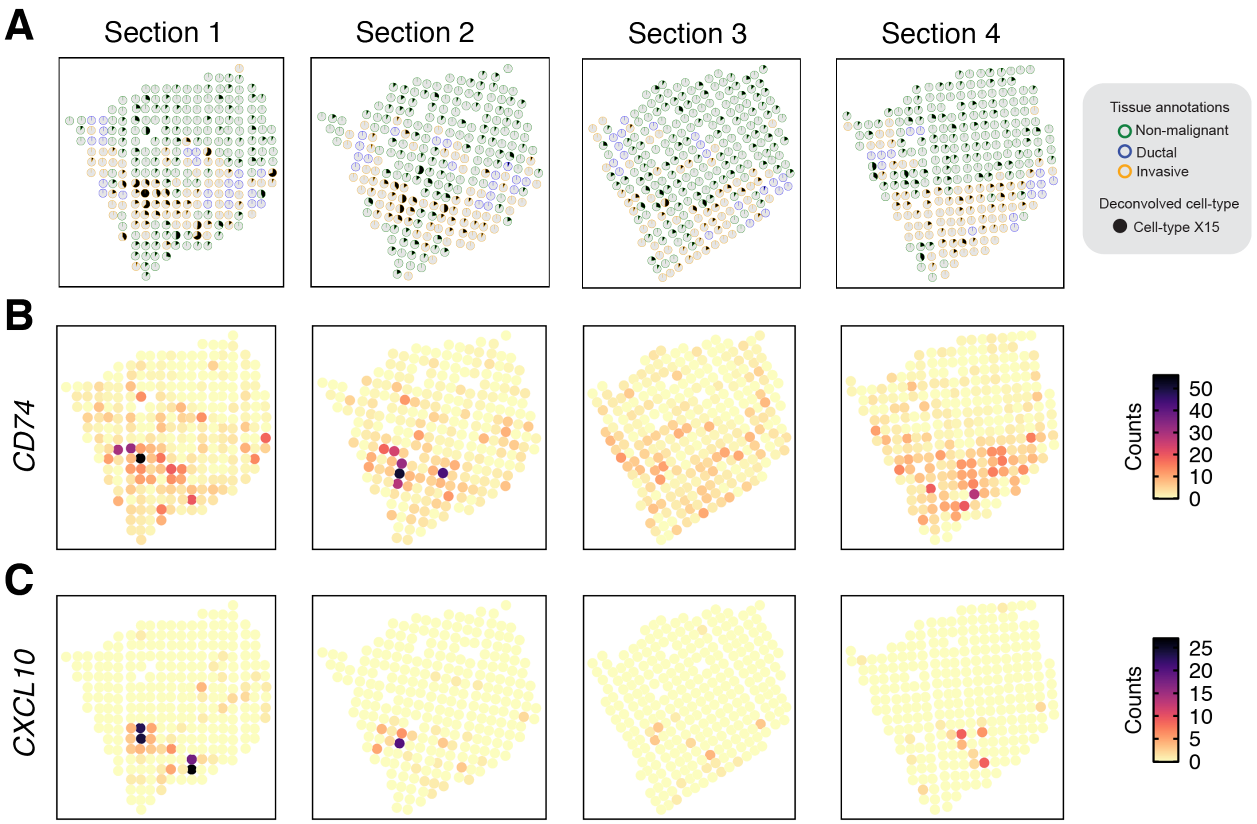


**Figure S18. STdeconvolve cell-types X15 of the breast cancer ST data.** A) Pixels of each breast cancer tissue section with outlines colored corresponding to their assigned tissue annotation. Pixels represent pie charts of cell-type pixel proportions, where the pixel proportion of deconvolved cell-type X15 is colored in black. B) Gene counts of top differentially upregulated genes for deconvolved cell-type X15.

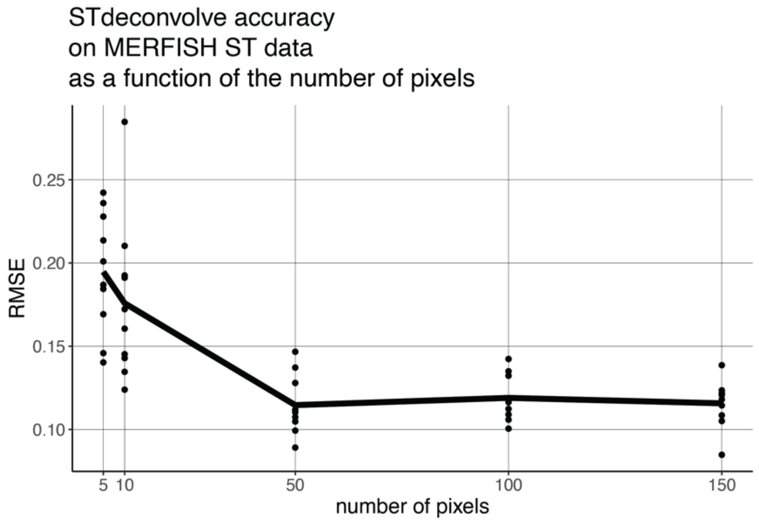


**Figure S19. Accuracy of deconvolution by STdeconvolve based on the number of pixels in the input dataset.** Each point is a replicate in which a random sample of pixels from the simulated MERFISH MPOA ST was used as the input into STdeconvolve. For each replicate, the mean RMSE was computed by averaging the RMSEs of the difference in deconvolved cell-type and ground truth cell-type proportions for each pixel. The solid line represents the mean RMSE of the replicates for each given number of pixels sampled.

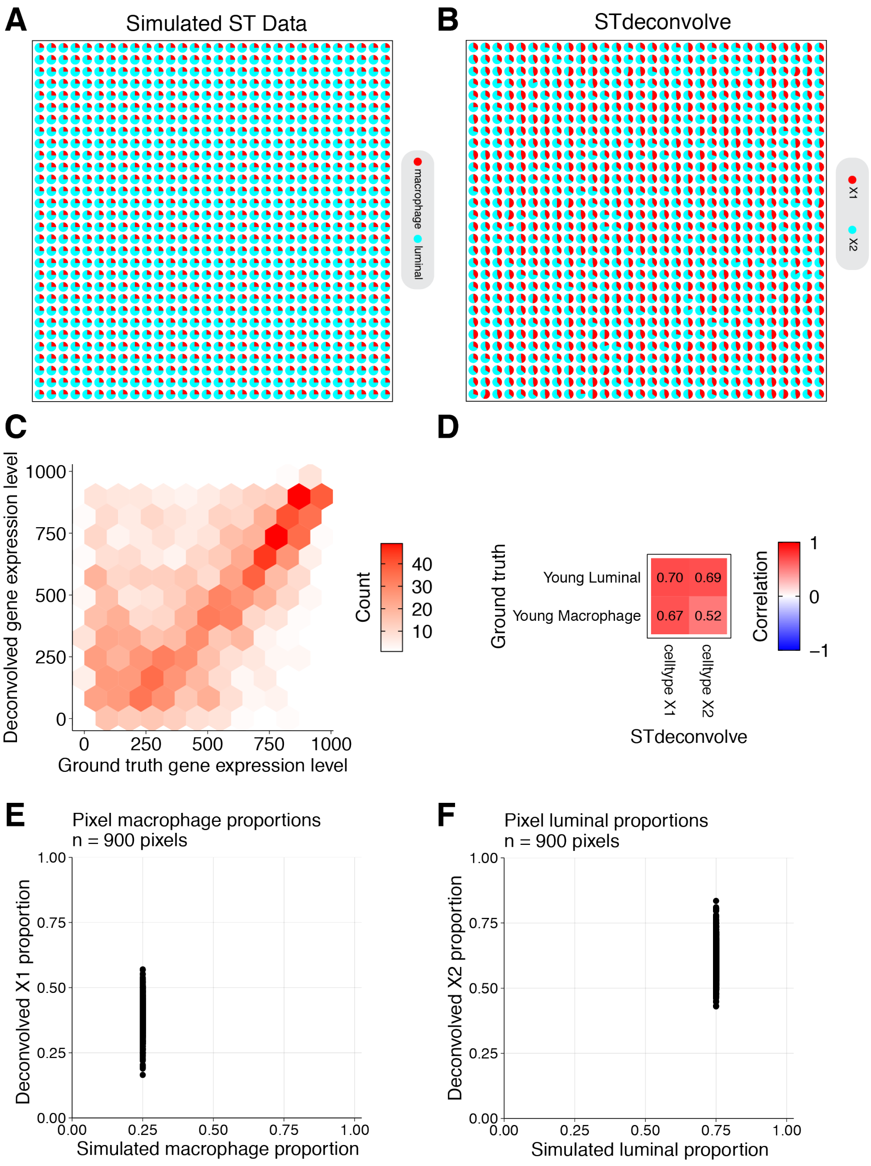


**Figure S20. Deconvolution failures.** A) Pie charts indicating the ground truth pixel proportions of luminal and macrophage cells with simulated uniform proportions across pixels. B) STdeconvolve predicted pixel proportions across pixels. C) The ranking of each gene based on its expression level in the transcriptional profiles of the deconvolved cell-types, compared to its gene rank in the transcriptional profile of the matched ground truth cell-types. D) Heatmap of Pearson's correlations between the transcriptional profiles of the ground truth cell-types and deconvolved cell-types. E-F) Observed ground truth pixel proportions compared to deconvolved pixel proportions for the macrophage and luminal cells.

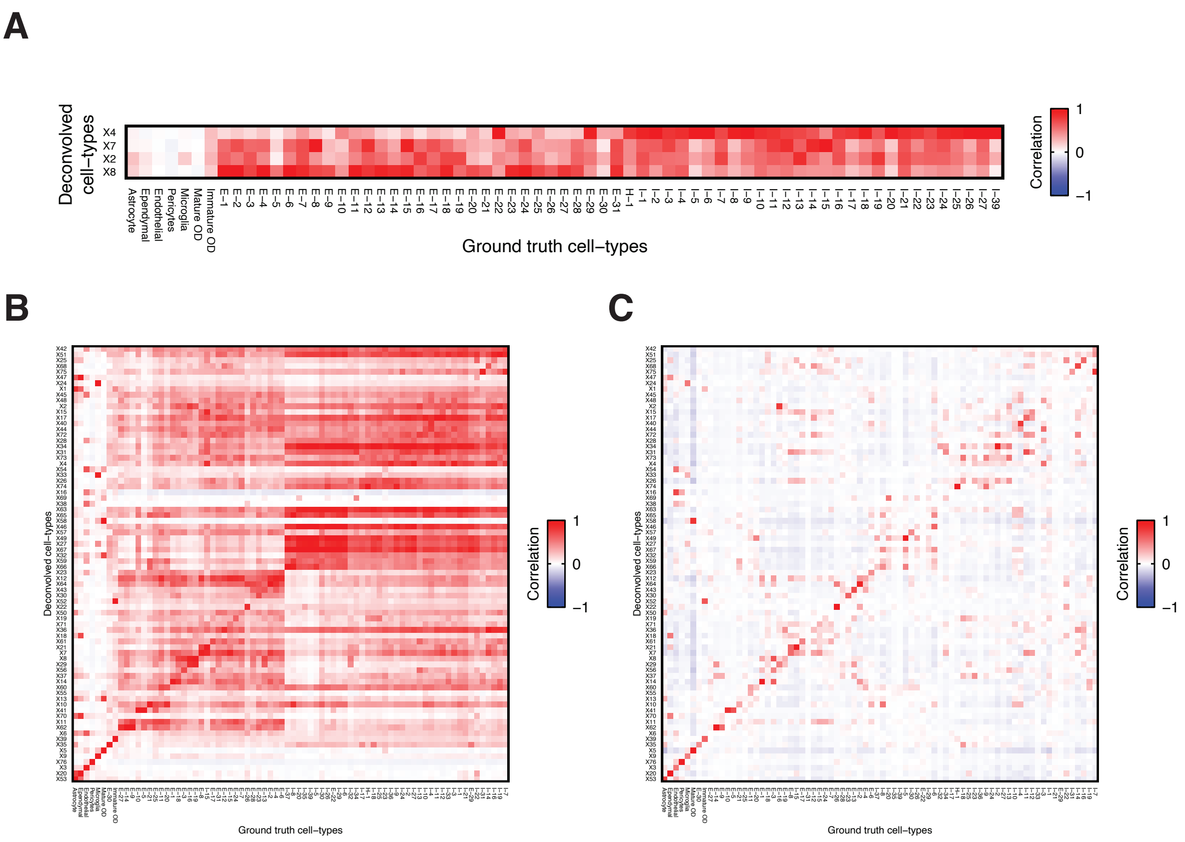


**Figure S21. Comparison of STdeconvolve cell-types to ground truth neuronal subtypes of simulated MERFISH MPOA ST data.** A) Heatmap of Pearson's correlations between the transcriptional profiles of the 76 ground truth cell-types and neuronal subtypes in the MERFISH MPOA data and STdeconvolve cell-types that matched to excitatory and inhibitory neuronal major cell-type types. B) Heatmap of Pearson's correlations between the transcriptional profiles of the 76 ground truth cell-types and neuronal subtypes in the MERFISH MPOA data and 76 deconvolved cell-types by STdeconvolve. C) Heatmap of Pearson's correlations between the cell-type pixel proportions of the 76 ground truth cell-types and neuronal subtypes in the MERFISH MPOA data and 76 deconvolved cell-types by STdeconvolve.

### D. Supplementary Tables

Table S1. Significant GO terms for STdeconvolve cell-type X15 derived from the breast cancer ST dataset. Table columns: “term” = GO term; “p.val” = *p*-value; “q.val” = multiple testing adjusted *p*-value; “sscore” = GO term enrichment score; “edge” = GO term edge score.
